# Heat stress-induced condensation of G3BP1 in perinuclear P-bodies in *C. elegans’* germline

**DOI:** 10.64898/2026.03.05.709961

**Authors:** Diya Zang, Yifan Jing, Xinya Huang, Yan Kuang, Jiewei Cheng, Wenkai Wang, Demin Xu, Chengming Zhu, Di Chen, Zhongying Zhao, Xuezhu Feng, Shouhong Guang

## Abstract

Stress granules (SGs) are essential subcellular assemblies that enable cells to adapt to environmental stress, and their dynamic assembly and disassembly are critical for maintaining cellular homeostasis and reproductive capacity. However, the function and regulation of stress granules in germline cells remain mysterious. In this study, we characterized the function of GTBP-1, the homolog of human G3BP1, in response to heat stress in *Caenorhabditis elegans* and elucidated its regulation in stress granule dynamics. *gtbp-1* mutants exhibit pronounced temperature-dependent biodirectional reproductive characteristics. At lower temperatures, their brood size is higher than that of wild-type animals, whereas at 25 °C they are completely sterile. GTBP-1 is diffusely distributed at normal culturing temperatures but forms perinuclear stress granules upon heat shock in the germline. While the NTF2 domain is essential for germline stress granule formation, other domains, including the IDR, RRM and RGG, are required for the recovery phase after heat shock. GTBP-1 stress granules colocalize with the P-bodies, but not with other germ granules. The depletion of P-bodies prohibited the perinuclear GTBP-1 stress granule formation. A combination of forward genetic screening together with RNAi-based candidate screening identified the P body components, the RNA helicase LAF-1, the SUMO protein SMO-1, and the mTOR pathway effector RSKS-1(S6K) as key regulators of GTBP-1 germline stress granule formation. Together, this work revealed that germline stress granules may be subjected to multiple layers of regulations and GTBP-1 may safeguards reproductive homeostasis under temperature stress by coordinating germline stress granule condensation.

**Author Summary:** Reproduction is particularly vulnerable to environmental stress, yet germ cells must remain functional to ensure fertility. How germ cells protect themselves under stressful conditions is still not well understood. In this study, we used the nematode Caenorhabditis elegans to examine how germ cells respond to heat stress, focusing on GTBP-1, a conserved protein involved in stress granule formation. We found that GTBP-1 plays contrasting roles depending on temperature: under normal conditions it restrains reproduction, whereas under heat stress it becomes essential for maintaining fertility. When animals are exposed to elevated temperature, GTBP-1 rapidly forms granules around the nuclei of germ cells, these granules assemble at sites occupied by processing bodies, another type of RNA-containing structure, revealing a close spatial relationship between two stress-responsive compartments. We also found that this process is regulated by conserved factors involved in RNA regulation and nutrient-responsive signaling. Together, our findings show how germ cells reorganize RNA–protein assemblies to preserve reproductive capacity under stressful conditions.

## Introduction

Organisms frequently encounter diverse environmental stresses, including temperature fluctuations, oxidative stress, and nutrient deprivation. To preserve cellular homeostasis, cells have evolved rapid and reversible response mechanisms, among which stress granules (SGs) represent one of the most critical stress-induced subcellular structures^1, 2^. Stress granules are primarily composed of translationally stalled mRNAs and associated proteins, and they assemble via liquid-liquid phase separation (LLPS)^3, 4^. By transiently sequestering mRNAs, stress granules modulate their translation, stability, and degradation, thereby dynamically regulating gene expression at the post-transcriptional level^5–7^. Notably, germ cells and early embryos are particularly vulnerable to environmental stresses, including heat stress, and the fidelity of reproductive processes relies heavily on precise intracellular homeostasis^8–10^.

The formation of stress granules is orchestrated by a complex network of protein-protein interactions. Beyond the classic translation initiation factor eIF2, several nucleating factors, including Ras GTPase-activating protein-binding protein 1 and 2 (G3BP1/2), ubiquitin-associated protein 2-like (UBAP2L), and T-cell-restricted intracellular antigen-1 (TIA-1), have been demonstrated to be crucial for stress granule assembly^11–13^. Mechanistic studies indicate that G3BP1/2 drive stress granule formation through RNA-induced conformational changes and self-aggregation, a process that depends on the synergistic interplay between the NTF2L domain and RNA-binding domain (RBD)^14–16^. During heat shock, the subsequent disassembly of stress granules is mediated by ubiquitination of G3BP1^17^, underscoring its central role in the dynamic remodeling of stress granules. In *C. elegans*, GTBP-1, the homolog of the human G3BP1, has been confirmed to participate in stress granule assembly^18, 19^, providing an ideal model for studying the dynamic regulation of stress granules and their connection to reproductive function. Nevertheless, it remains unclear whether GTBP-1 directly governs germ cell homeostasis under temperature stress and how its activity is modulated by environmental conditions.

Germ granules are RNA-enriched condensates typically localized to the perinuclear region of germ cells, involved in RNA homeostasis and post-transcriptional regulation, and are crucial for germline development and reproductive homeostasis^20–22^. In *C. elegans*, germ granules comprise multiple substructures, including P granules, Z granules, Mutator foci, SIMR foci, D granules, E granules and P-bodies ^21, 23–25^. In yeast and mammalian cells, stress granules and P-bodies exhibit partial spatial overlap and undergo dynamic messenger ribonucleoprotein (mRNP) exchange under stress conditions; with some components being targeted for autophagic degradation^26–28^. These observations suggest that the material flux and functional interactions among diverse RNA-associated condensates form an integrated network that is critical for cellular stress resilience.

This study aims to investigate the role of GTBP-1 in regulating reproductive fitness under temperature variations and to elucidate how stress granules are regulated in the germline in *C. elegans*. Through a combination of genetic approaches, cell biological assays, and RNAi-based screening, we characterize the domain-specific functions of GTBP-1, its interactions with other stress granule components, and its spatial organization within germ granules. We further assessed the regulatory influence of RSKS-1 on GTBP-1 stress granule assembly and reproductive homeostasis. Together, our findings provide new mechanistic insights into how temperature stress modulates germline stress granule formation and the maintenance of reproductive homeostasis.

## Results

### Heat shock–induced perinuclear stress granule formation in *C. elegans’* germline

*C. elegans’* GTBP-1 is a homolog of human G3BP1, which is the core factor of stress granules. GTBP-1 contains a Nuclear Transport Factor 2 (NTF2) domain, an internal disorder region (IDR) domain, a RRM and an Arginine/Glycine-rich (RGG) domain. The NTF2 domain is reported to engage in mediating macromolecular shuttling between the nucleus and the cytoplasm. The IDR domain may participate in liquid-liquid phase separation. Both RRM and RGG domains are likely involved in binding RNAs. To elucidate the functions of GTBP-1 in germline, we first generated full-length *gtbp-1* knockout alleles (*ust600* and *ust601*) using CRISPR/Cas9 technology (Figure 1A). The *gtbp-1(ust600)* deleted most of the amino acid sequences and *gtbp-1(ust601)* deleted nearly the whole gene sequence, which is likely a null allele. Both mutants exhibited no overt developmental defects, but they showed pronounced reproductive abnormalities upon temperature alternations. At 15 °C and 20 °C, *gtbp-1* mutants produced significantly more progeny than N2 animals (Figure 1B), whereas at 25 °C they became sterile, suggesting that GTBP-1 is essential for maintaining reproductive homeostasis under different temperature conditions.

**Figure 1.**
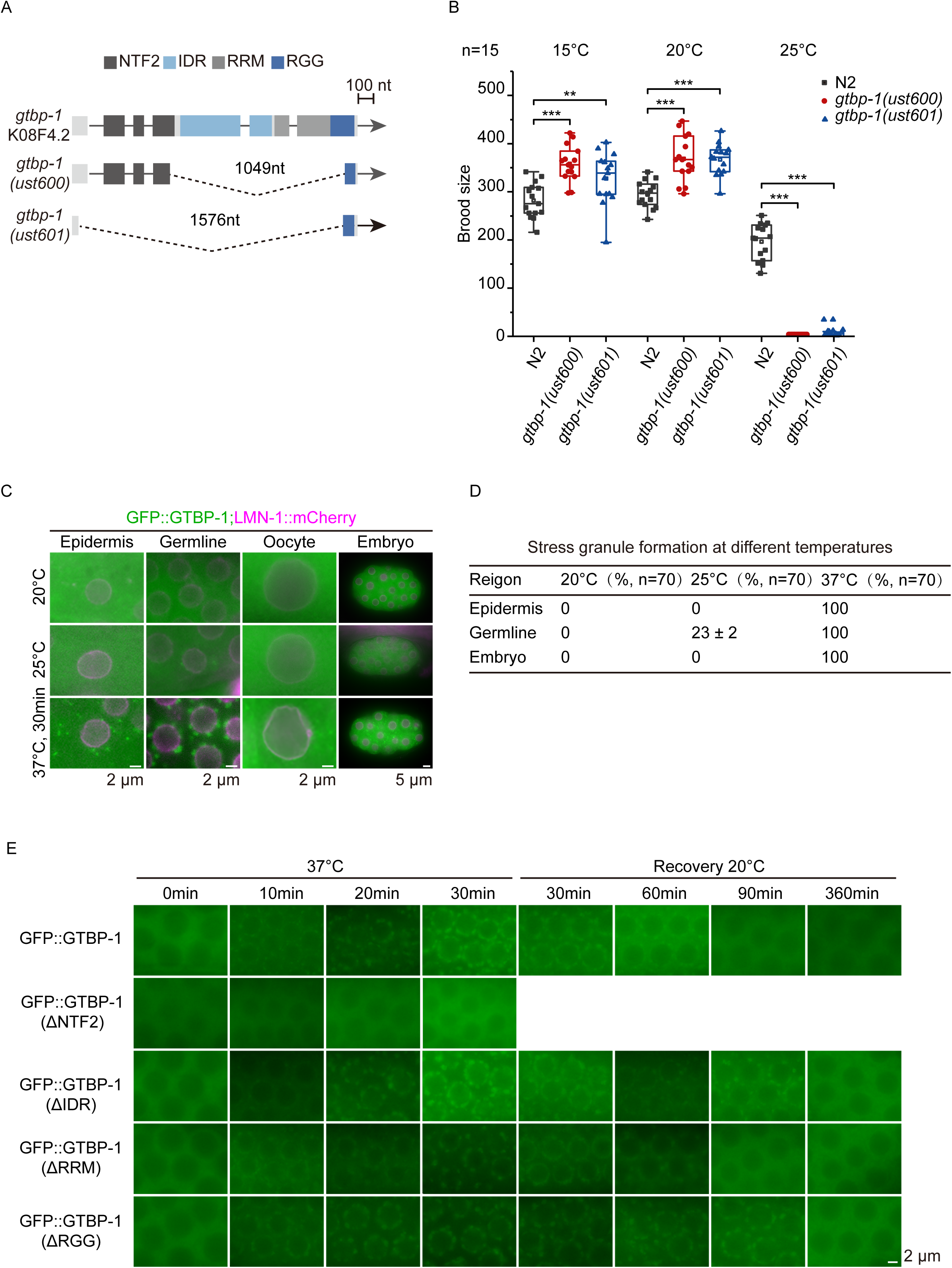
GTBP-1 mediates reproductive homeostasis and temperature-dependent stress granule assembly. (A) Schematic representation of the *gtbp-1* gene structure and CRISPR/Cas9-generated deletion alleles (*ust600, ust601*). Domains: Nuclear Transport Factor 2 (NTF2) domain, internal disorder region (IDR) domain, RRM and Arginine/Glycine-rich (RGG) domains. (B) Brood size analysis of *gtbp-1* mutants at 15 °C, 20 °C, and 25 °C, respectively (n=15 animals). Box plots show median, 25–75% percentiles, and minimum/maximum values. (C) Representative images of GFP::GTBP-1 localization in multiple tissues under different temperature conditions. GTBP-1 displays diffuse cytoplasmic distribution at 20 °C but relocalizes into perinuclear granules upon heat stress (25 °C chronic exposure or 37 °C acute heat shock for 30 min). LMN-1::mCherry was used to label the nuclear envelope. Images are representative of at least three animals per condition. (D) Quantification of stress granule formation across tissues and temperatures. Stress granule assembly is enhanced at 25 °C in a tissue-specific manner and is strongly induced by acute heat shock (n=70 animals). (E) Domain analysis of GTBP-1. Deletion of the NTF2 domain abolishes stress granule assembly, whereas IDR or RGG deletion permits assembly but delays disassembly during recovery. Images are representative of at least three animals per condition. Data are presented as mean ± SEM from three independent experiments. Statistical significance was determined by a two-tailed t-test. *p < 0.05; **p < 0.01; *p < 0.001; ns, not significant (p > 0.05).

We generated GFP tagged GTBP-1 and examined GFP::GTBP-1 distribution across multiple tissues and monitored stress granule assembly under different temperature conditions (Figure 1C–D). The nuclear envelope was labeled by the lamin protein LMN-1::mCherry. Under normal lab culturing conditions at 20 °C, GFP::GTBP-1 exhibited an evenly cytoplasmic distribution in epidermal cells, germ cells, and early embryos, with no detectable stress granule assembly. When the temperature increased to 25 °C, small GTBP-1 granules emerged specifically in the germline cells, whereas no granules were detected in other tissue, indicating that chronic 25 °C culture is sufficient to trigger stress granule condensation. Upon acute heat shock at 37 °C for 30 minutes, GTBP-1 formed robust stress granules across multiple tissues. Especially, we noticed that GTBP-1 accumulated prominently in the perinuclear granules in germline cells.

To dissect the domain-specific contributions of GTBP-1 to germline stress granule assembly, we generated transgenic strains expressing GFP::GTBP-1 variants lacking individual protein domains and examined stress granule formation in the germline upon heat shock (Figure 1E). The deletion of the NTF2 domain completely abolished stress granule formation, which is consistent with the findings for mammalian G3BP proteins^14, 29^ and implies that the nuclear and cytoplasmic shuttling of GTBP-1 might be indispensable for stress granule formation. In contrast, the deletion of the IDR or RGG domain did not prevent stress granule assembly; however, stress granule disassembly during recovery from heat shock was markedly delayed, suggesting that these domains might primarily modulate stress granule fluidity and dissolution rather than heat shock–induced nucleation. Together, these results revealed that heat shock could induce perinuclear stress granule formation of GTBP-1 in *C. elegans’* germline.

### GTBP-1 colocalized with PQN-59 and TIAR-1 within perinuclear stress granules in the germline

PQN-59 and TIAR-1 are two key stress granule proteins reported to involve in stress granule formation^13, 18^. PQN-59 is an ortholog of human UBAP2 (ubiquitin associated protein 2) and UBAP2L (ubiquitin associated protein 2 like)^18, 30^. TIAR-1 is a homolog of the mammalian TIA-1/TIAR family of proteins, which contains a prion-like domain, exhibits mRNA binding ability and is reported to accumulate in germline P granules^31^. We tested their colocalization with the GTBP-1 stress granules upon heat shock. At the normal laboratory cultivation temperature at 20 °C, PQN-59 is evenly distributed in the cytoplasm in epidermis, germline cells, and early and late embryos. Upon heat shock at 37 °C for 30 min, PQN-59 accumulated in perinuclear stress granules and extensively overlapped with GFP::GTBP-1 signals (Figure 2A). In epidermis and early and late embryos, GTBP-1 and PQN-59 granules colocalized in the cytoplasm.

**Figure 2.**
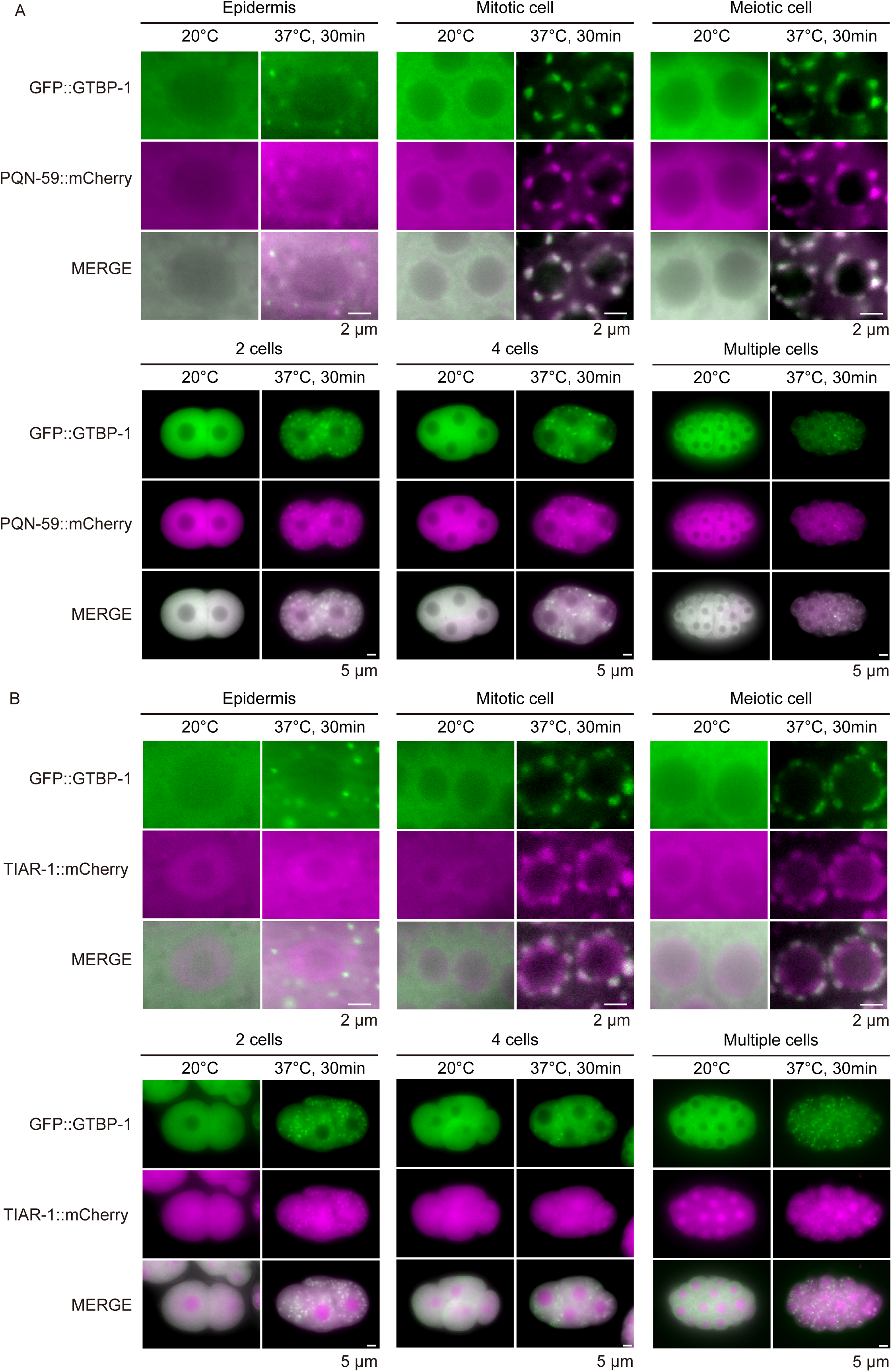
GTBP-1 colocalizes with PQN-59 and TIAR-1. (A) Fluorescence images of animals expressing GFP::GTBP-1 (green) and PQN-59::mCherry (magenta) in epidermal cells, mitotic germ cells, meiotic germ cells and early embryos (2-cell, 4-cell, and multicellular stages) at 20 °C and following heat shock (37 °C for 30 min). (B) Fluorescence images of animals expressing GFP::GTBP-1 (green) and TIAR-1::mCherry (magenta) in epidermal cells, mitotic germ cells, meiotic germ cells and early embryos (2-cell, 4-cell, and multicellular stages) at 20 °C and following heat shock (37 °C for 30 min). Images are representative of at least three animals per condition.

In contrast, at 20 °C, TIAR-1 was distributed in both the cytoplasm and nucleoplasm (Figure 2B). Upon 37 °C heat shock, in embryos and epidermis, TIAR-1, but not GTBP-1, was enriched in the nucleus. In the germline, although TIAR-1 and GTBP-1 were both recruited to perinuclear germ granules upon heat stress, they displayed clear intra-granular spatial segregation. TIAR-1 and GTBP-1 occupied overlapping yet distinct subregions within the same perinuclear granules, rather than being uniformly mixed, indicating internal spatial heterogeneity within germline stress granules.

Collectively, these results suggested that GTBP-1 can co-aggregate with different stress granule proteins, yet exhibiting cell type specificity.

### RNAi-based candidate screening identifies LAF-1 and SMO-1 that are required for GTBP-1 germline stress granule formation

Notably, although PQN-59 and TIAR-1 colocalized with GTBP-1 upon heat shock, knockdown of PQN-59 or TIAR-1 by RNAi has minimal, if any, effect on GTBP-1–associated stress granule formation in both embryos and germline upon 37 °C heat shock (Figure 3A–B), suggesting that both PQN-59 and TIAR-1 are dispensable for heat shock–induced GTBP-1 granule assembly in the germline. Interestingly, at normal laboratory culture conditions at 20 °C, upon knockdown PQN-59 by RNAi, GTBP-1 protein aggregated and formed perinuclear granule structures in Z2/Z3 cells, the two germ cells, in embryos.

**Figure 3.**
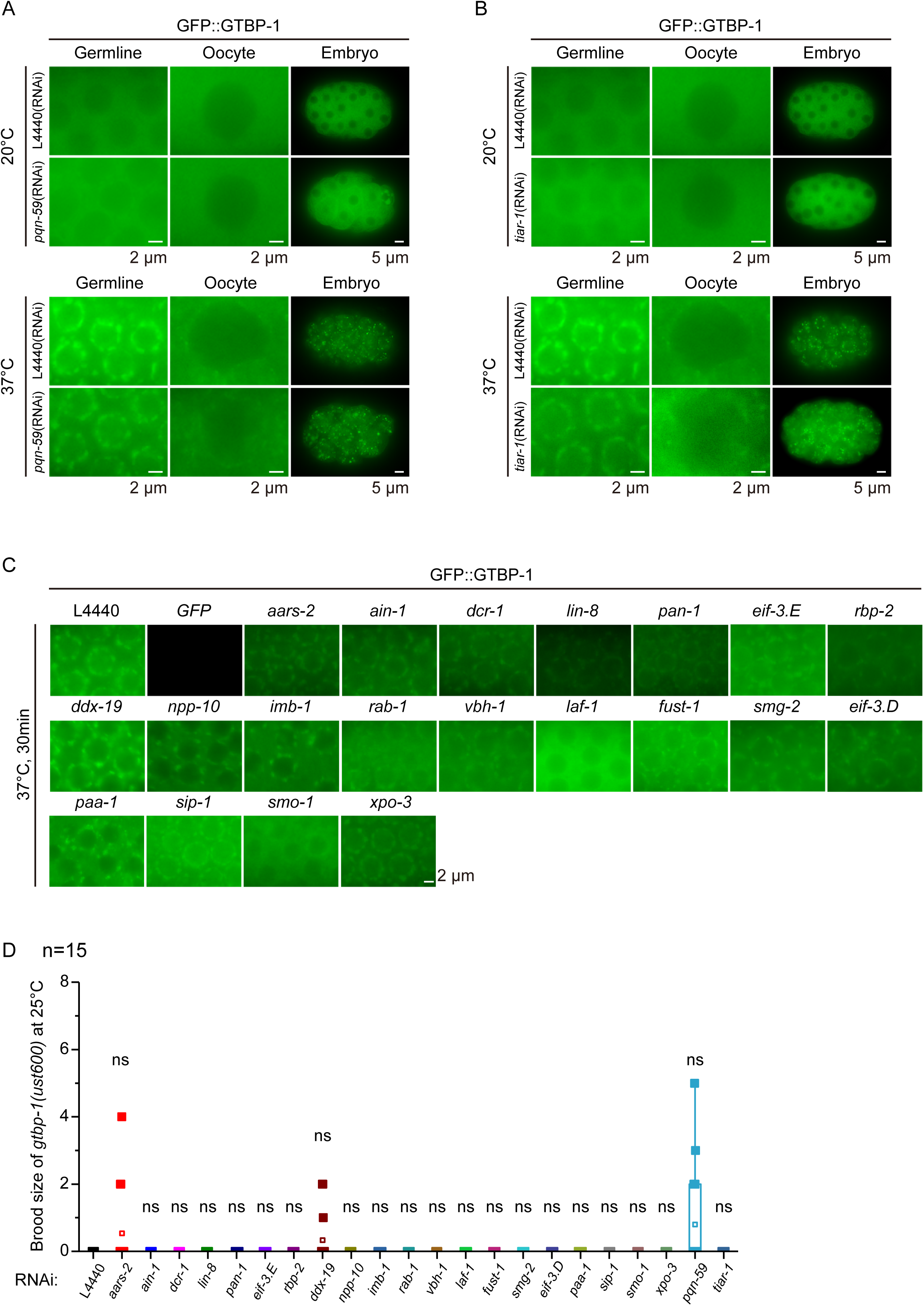
RNAi screening identifies PQN-59, LAF-1, and SMO-1 as regulators of GTBP-1–associated stress granule formation. (A–B) Fluorescence images of animals expressing GFP::GTBP-1 (green) in the germline, oocytes, and embryos under RNAi conditions at 20 °C (upper panels) or heat shock at 37 °C for 30 min (lower panels). (A) Effect of *pqn-59*(RNAi). (B) Effect of *tiar-1*(RNAi). Images are representative of at least three animals per condition. (C) RNAi screen of multiple candidate genes showing their effects on GFP::GTBP-1 granules in the germline under heat shock (37 °C for 30 min). (D) Brood size analysis of *gtbp-1* mutants at 25 °C following RNAi knockdown of candidate genes (n=15 animals). Box plots show median, 25–75% percentiles, and minimum/maximum values. Data represent results from three independent experiments. Statistical significance was determined by a two-tailed t-test. *p < 0.05; **p < 0.01; *p < 0.001; ns, not significant (p > 0.05).

We searched for the factors required for GTBP-1 stress granule formation in the germline by feeding RNAi targeting both known core components of stress granules and predicted GTBP-1-interacting proteins^14, 29, 32, 33^. We knocked 35 genes individually by RNAi. The knockdown of *laf-1* and *smo-1* significantly abolished heat shock–induced GTBP-1–associated stress granule formation (Figure 3C). LAF-1 is a DDX3 class RNA helicase that accumulates in both cytoplasm and perinuclear germline P granules, and is involved in asymmetric phase separation and germ cell fate determination^23, 34, 35^. SMO-1 is the ortholog of human SUMO1 (small ubiquitin like modifier 1), suggesting the regulation of germline stress granule by SUMOylation. However, the depletion of these genes at 25 °C by RNAi did not rescue the reproductive defects of *gtbp-1* mutants (Figure 3D). Collectively, these results revealed that LAF-1 and SMO-1 serve as key regulators controlling GTBP-1 germline stress granule condensation.

### GTBP-1 colocalized with P-bodies in germ cells and early embryos

Our recent work has revealed 7 perinuclear germ granules in *C. elegans’* germline cells. To assess the spatial relationship of heat shock–induced GTBP-1 stress granule with these germ granules, we crossed GFP::GTBP-1 with the seven tagRFP or mCherry-tagged germ granule strains (Figure 4A). Among the marker proteins, PGL-1 stands for P granules, CGH-1 for P-bodies, ZNFX-1 for Z granules, MUT-16 for Mutator foci, SIMR-1 for SIMR foci, ELLI-1 for E granules and DDX-19 for D granules ^24, 25, 36–39^.

**Figure 4.**
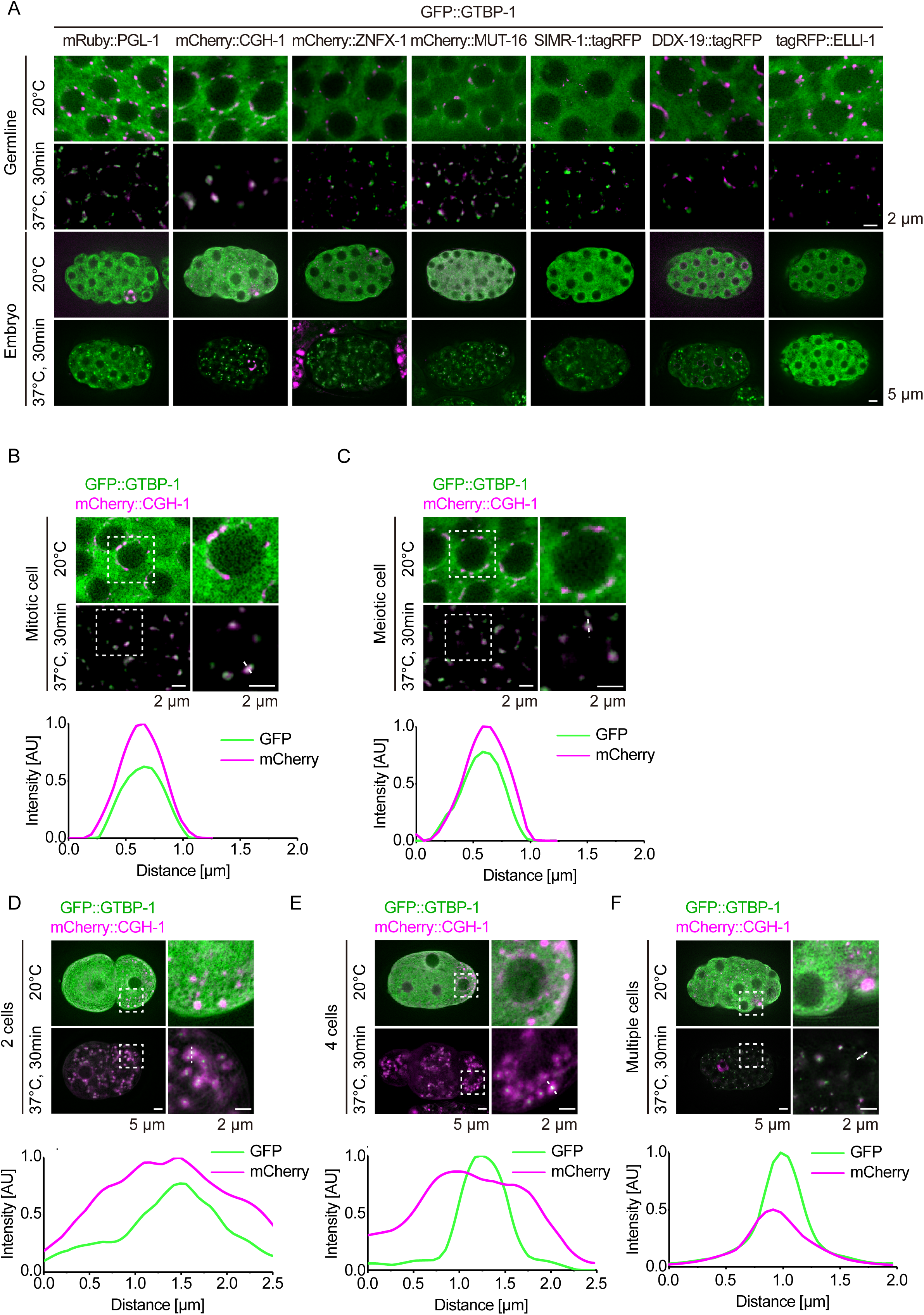
GTBP-1-associated stress granules colocalized with perinuclear P-bodies in the germline. (A) Subcellular colocalization assay of GFP::GTBP-1 with seven germ granule markers (mRuby::PGL-1 for P granule, mCherry::CGH-1 for P-body, mCherry::ZNFX-1 for Z granule, mCherry::MUT-16 for mutator foci, SIMR-1::tagRFP for SIMR granule, DDX-19::tagRFP for D granule, and tagRFP::ELLI-1 for E granule) in the germline and early embryos at 20 °C or following heat shock (37 °C for 30LJmin). (B–F) Colocalization of GFP::GTBP-1 (green) with mCherry::CGH-1 (magenta) in different tissues and developmental stages: (B) Mitotic cells of the germline, (C) Meiotic cells of the germline, (D) Two-cell embryo, (E) Four-cell embryo, and (F) Multicellular embryo. Fluorescence intensity along the dotted line was quantified using ImageJ. All images are representative of more than three animals.

Upon heat shock at 37 °C for 30 min, GTBP-1–associated stress granules showed extensive overlap with the P-body marker mCherry::CGH-1, whereas they exhibited only partial overlap with P granules and Mutator foci (Figure 4A–C) in the germline. In embryos, heat shock–induced GTBP-1 granules also displayed colocalization with CGH-1 (Figure 4D–F).

We then tested whether P-body per se is required for the heat shock–induced GTBP-1 stress granule formation. We knocked down the core P-body components, including *cgh-1*, *edc-3* and *ifet-1*, which encode conserved factors involved in P-body assembly and mRNA metabolism, by feeding RNAi (Figure 5)^40, 41^. Consistent with their roles in P-body formation, knocked down *edc-3* and *ifet-1* disrupted the mCherry::CGH-1-marked perinuclear P-body assemblage in germline. And knock down *cgh-1, edc-3* or *ifet-1* prohibited the formation of heat shock–induced condensation of GTBP-1 to the perinuclear P-body.

**Figure 5.**
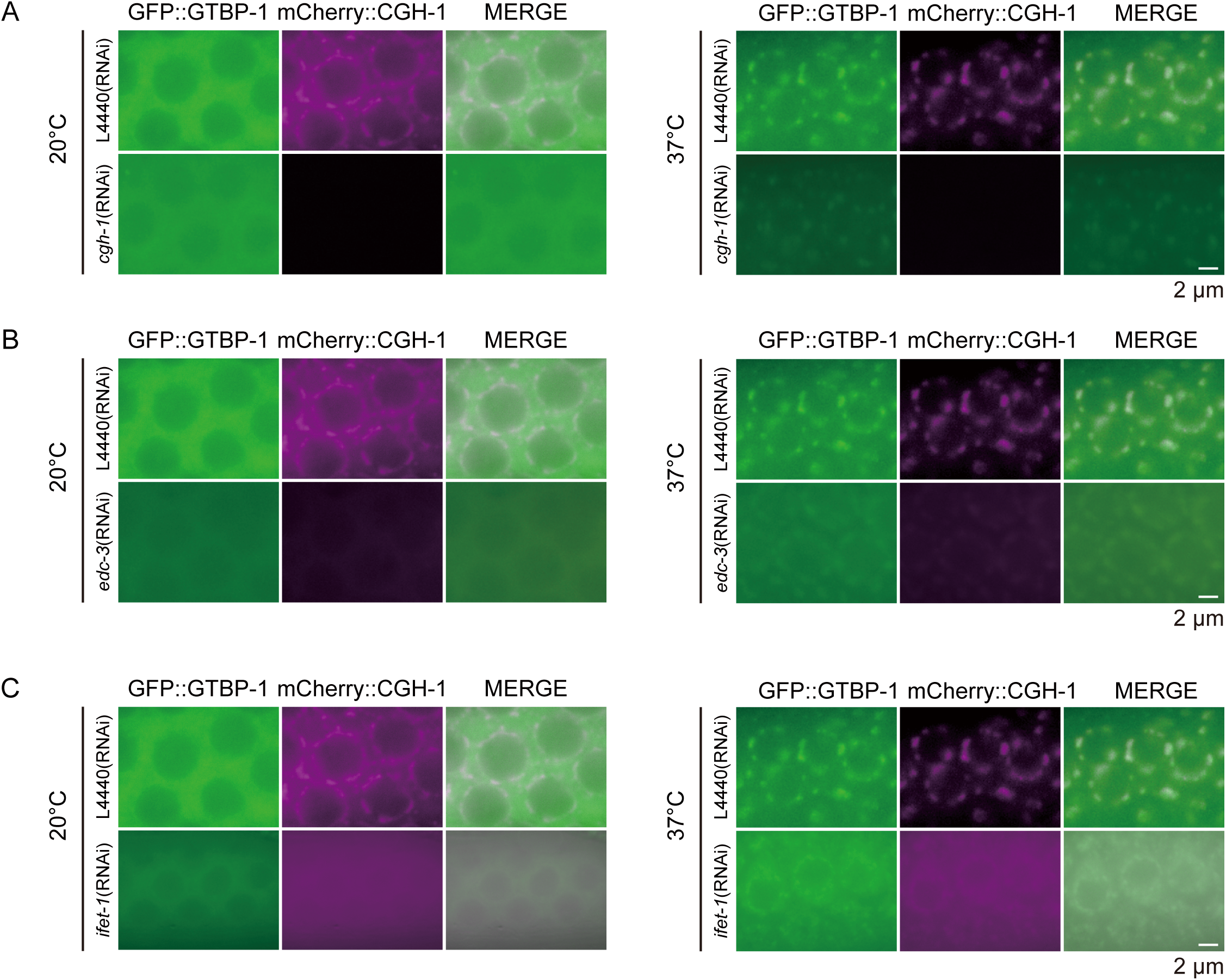
Disruption of P-body components abolished GTBP-1 stress granule assembly. (A–C) Representative fluorescence images showing GFP::GTBP-1 and mCherry::CGH-1 subcellular localization in young adult and gravid adult germlines under RNAi against *cgh-1* (A), *edc-3* (B), or *ifet-1* (C) at 20 °C and after 37 °C heat shock for 30 min. Images are representative of at least three animals per condition.

Thus, these data suggested that heat shock could induce the recruitment of GTBP-1 stress granules to the perinuclear P-bodies in germline cells.

### Forward genetic screening identified RSKS-1 modulates GTBP-1–dependent fertility and stress granule assembly

To investigate the biological roles of heat shock–induced perinuclear GTBP-1 stress granule formation, we performed a forward genetic screen to search for mutants that rescue the sterility of *gtbp-1* mutants at 25 °C. *gtbp-1(ust600)* animals were chemically mutagenized and F2 animals that are fertile at 25 °C were selected. We isolated a *rsks-1*(*ust761*) mutant that restored the fertility of the *gtbp-1* mutant (*ust600*) from approximately 20,000 haploid genome (Figure 6A–B). *rsks-1*(*ust761*) mutant changed an amino acid Histidine200 to Asparagine. Then we acquired a *rsks-1(ok1255)* allele from CGC and also generated another allele *rsks-1(ust763)* by CRISPR-Cas9 technology. All these *rsks-1* alleles, as well as knockdown *rsks-1* by feeding RNAi, modestly rescued the fertility of *gtbp-1* mutants at 25 °C (Figure 6B).

**Figure 6.**
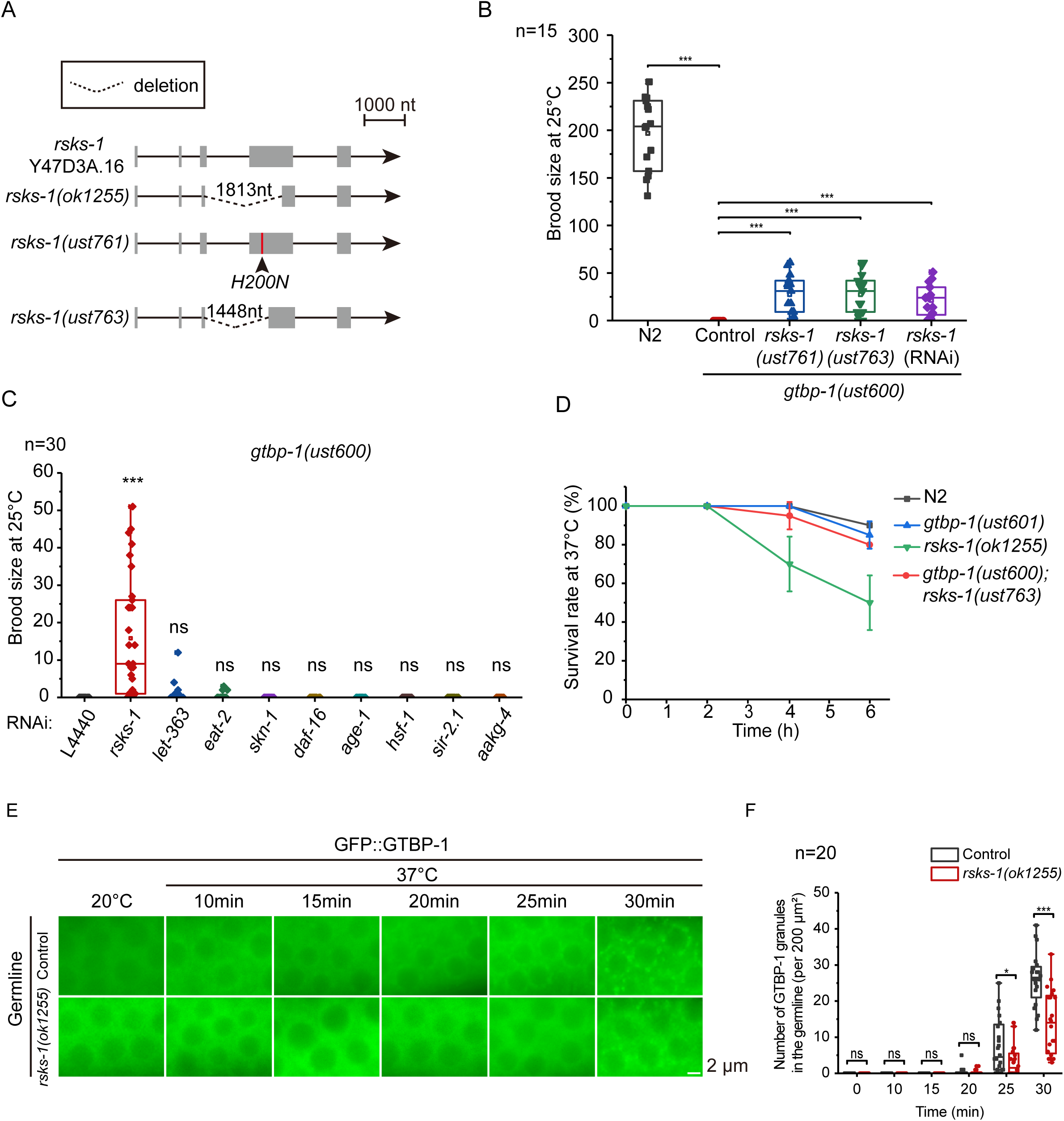
RSKS-1 modulates GTBP-1–dependent reproductive homeostasis and stress granule assembly. (A) Schematic representation of the *rsks-1* gene structure and the alleles (*ok1255, ust761, ust763*) generated by CRISPR-Cas9 and chemical mutagenesis. (B) Brood size analysis of N2, *gtbp-1* mutant, and *rsks-1;gtbp-1* double mutants at 25 °C (n=15 animals). Box plots show median, 25–75% percentiles, and minimum/maximum values. (C) Brood size analysis of *gtbp-1* mutants at 25 °C following RNAi knockdown of selected lifespan-regulating genes (n=15 animals). Box plots show median, 25–75% percentiles, and minimum/maximum values. (D) Survival curves of various experimental groups at 37 °C. Data represent mean ± SEM from three independent experiments; error bars indicate SEM. (E) Expression of GFP::GTBP-1 in the germline of wild-type (control) and *rksk-1(ok1255)* worms following 37 °C heat shock. All images are representative of more than three animals. (F) Quantification of GFP::GTBP-1 stress granules in the germline under the same conditions as in (E). Granules were counted using fixed ROIs of approximately 200 μm², each dot representing one unit area, sampled from ≥10 animals. Box plots show median, 25–75% percentiles, and minimum/maximum values. All images are representative of more than three animals. Statistical significance was determined by a two-tailed t-test. *p < 0.05; **p < 0.01; *p < 0.001; ns, not significant (p > 0.05).

In mammalian cells, S6K localizes to oxidative stress–induced stress granules and contributes to their assembly and maintenance^42^. RSKS-1, the *C. elegans* homolog of S6K and a key effector of the mTOR pathway, is known to regulate translation and lifespan^43^. To test whether the suppression of *gtbp-1* sterility results from a general reduction in lifespan-regulatory or mTOR signaling, we used feeding RNAi to target multiple well-characterized longevity genes^44–46^, yet knock down none of these genes could rescue the fertility of *gtbp-1* mutant at 25 °C (Figure 6C), suggesting a specific effect of RSKS-1 in modulating the fertility of *gtbp-1* animals.

Heat-shock survival assays revealed a similar reduction of survival between *gtbp-1* and wild type N2 animals. Interestingly, the mutation of *rsks-1* further promoted the heat shock–induced lethality than wild type N2 animals (Figure 6D). The *rsks-1; gtbp-1* double mutant revealed similar survival to that of wild type N2 animals upon heat shock, suggesting that RSKS-1 and GTBP-1 may counteract each other in mediating stress reactions.

Strikingly, RSKS-1 promotes the heat shocked-induced GTBP-1 germline stress granule aggregation. We examined the assembly dynamics of GTBP-1 stress granules under heat shock. In the absence of *rsks-1*, the formation of GTBP-1 stress granules was markedly delayed, and the number of granules was reduced (Figure 6E–F).

To further delineate how RSKS-1 is involved in heat stress responses, we characterized the subcellular distribution of RSKS-1 upon heat stress. LMN::mCherry was used to display the nuclear membrane. Under normal laboratory culturing temperature at 20 °C, RSKS-1 is distributed in the cytoplasm and nucleoplasm. Upon exposure to 37 °C for 30 min, RSKS-1 underwent a striking relocalization and accumulated in discrete perinuclear granule-like structures in germline and oocyte. In embryos, RSKS-1 was enriched in the nucleus and formed puncta in the cytoplasm (Figure 7A). These heat-induced RSKS-1 perinuclear granules showed substantial colocalization with the P-body marker CGH-1 in germ granule (Figure 7B).

**Figure 7.**
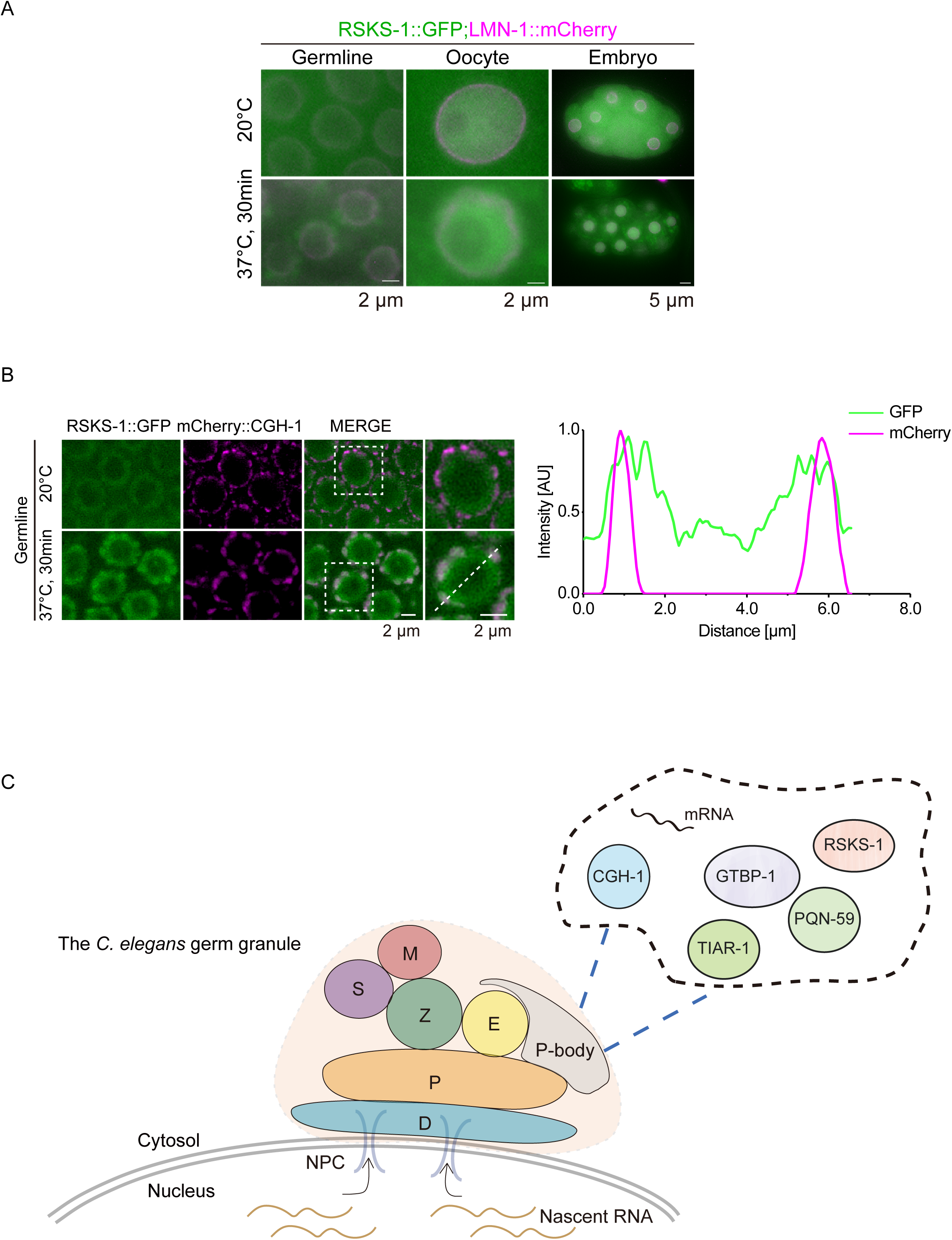
Heat stress induces perinuclear accumulation of RSKS-1 in P-body. (A) Representative images of RSKS-1::GFP localization in multiple tissues at 20 °C and following heat shock (37 °C for 30 min). LMN-1::mCherry was used to label the nuclear envelope. (B) Representative images of the subcellular localization of RSKS-1::GFP (green) and the P-body marker mCherry::CGH-1 (magenta) in the germline under culture conditions (20 °C) or heat stress (37 °C for 30 min). Fluorescence intensity along the dotted line was quantified using ImageJ. (C) A working model of heat shock–induced GTBP-1 stress granule condensation in the perinuclear P-bodies. The *C. elegan’s* germline contains approximately 7 perinuclear germ granules, among which the P-body is marked by CGH-1, EDC-3 and IFET-1. Upon heat stress, P-bodies enrich PQN-59, TIAR-1, RSKS-1, and GTBP-1 to the perinuclear germline granules.

Together, these results identify RSKS-1 as a heat-responsive regulator that promotes GTBP-1 stress granule assembly and modulates GTBP-1–dependent fertility under thermal stress.

## Discussion

Stress granules have been widely recognized as key regulators of cellular stress responses, yet their germline-specific assembly and physiological significance remain poorly understood. Here, using *C. elegans*, we identify GTBP-1 as a principal regulator of temperature-dependent reproductive homeostasis. Upon heat stress, GTBP-1 is recruited to perinuclear P-bodies in germ cells. PQN-59, LAF-1, SMO-1 and RSKS-1, as well as the P-body components, are key regulators of GTBP-1 germline stress granule formation (Figure 7C). We speculated that GTBP-1–containing condensates act as key stress-responsive structures required for maintaining germline homeostasis and fecundity under adverse environmental conditions.

An interesting finding is that *gtbp-1* mutant animals produce more progeny at low temperatures at 15 °C and 20 °C, but become completely sterile at higher temperatures at 25 °C, establishing GTBP-1 as a temperature-responsive factor essential for germline integrity. These results suggested that GTBP-1 may normally restrain reproductive output under favorable conditions but is required to maintain reproductive homeostasis under adverse environmental conditions. Consistently, mammalian G3BP proteins exert context-dependent, bidirectional regulation of cell proliferation^8, 11, 47, 48^. This dual role indicates that GTBP-1 not only serves as a core component of stress granules but also fine-tunes germline physiology to maintain reproductive capacity.

Domain analysis reveals that the loss of the NTF2 domain fully abrogates stress granule formation, mirroring the essential nucleating role of NTF2L in mammalian G3BP1/2^14, 15^. In contrast, deletion of the IDR or RGG region preserves initial granule assembly but impairs disassembly during recovery, highlighting their contribution to stress granule fluidity and reversibility^49, 50^. These observations suggested that GTBP-1–associated stress granules may be built through a domain-encoded phase-separation mechanism and that dynamic turnover, rather than merely granule formation, is crucial for preserving germline homeostasis under stress. The internal organization of GTBP-1–associated stress granules further support the view that stress granules are structured, multifunctional condensates. GTBP-1 extensively overlaps with PQN-59 but forms spatially distinct subregions relative to TIAR-1, consistent with a core–shell architecture reported in other systems^4, 51, 52^.

RNAi-based candidate screening identified that LAF-1, PQN-59, and SMO-1 are required for GTBP-1 condensation. The depletion of LAF-1 strongly suppresses stress granule assembly, consistent with its activity as an RNA helicase that promotes RNA–protein granule nucleation^34, 35, 53^. In *C. elegans*, *smo-1* encodes the sole SUMO protein, and SUMOylation is elevated under stress^54–56^. Knock down *smo-1* expression markedly diminishes GTBP-1 condensation during heat stress, suggesting that SUMOylation modulates the phase-separation properties of GTBP-1 or its partners and thereby contributes to stress granule formation. However, neither factor rescues the sterility of *gtbp-1* mutants at 25 °C, underscoring that GTBP-1 performs additional germline-specific functions.

Our study reveals an intimated relation between GTBP-1–contained stress granules and the P-body in the *C. elegans* germline under heat stress. GTBP-1-assocciated stress granule colocalized with the P-body, but not other known perinuclear germ granules which are key players in small RNA and RNAi related processes. Depletion of core P-body components, including *cgh-1, edc-3* and *ifet-1*, disrupted the P-body formation in perinuclear regions and therefore abrogated the heat shock–induced GTBP-1 germline granule formation. The dependency of GTBP-1 stress granules on heat stress and the presence of P-body suggested that the stress granules are likely dynamic structures capable of exchanging components, functionally complementing, or interacting RNAs, therefore likely contributing to protein and RNA homeostasis in germ cells to safeguard reproductive integrity under thermal stress^57–59^. Whether GTBP-1 per se or P-body factors are sumoylated to facilitate their interplay upon heat stress require further investigation.

The mechanistic target of rapamycin (mTOR) signaling pathway plays a central role in multicellular organisms by coordinating protein translation, energy metabolism, and lifespan regulation^60–62^. Its downstream effector, ribosomal S6 kinase (S6K), has been shown in mammals to participate in stress granule assembly under oxidative stress. Notably, loss of the *C. elegans* homolog, RSKS-1, attenuates heat stress-induced germline stress granule formation and markedly decreases worm survival^42^. Nevertheless, whether and how RSKS-1 responses to heat stress and consequently impacts cellular homeostasis and reproductive function remains mysterious. Here, we found that the loss of *rsks-1* partially rescued the sterility of *gtbp-1* mutant at 25 °C. Upon heat stress, RSKS-1 undergoes a pronounced relocalization and forms perinuclear granule-like structures, and further recruits GTBP-1 to P-body, suggesting that RSKS-1 not only acts as a global metabolic regulator but also as a stress granule-embedded factor that may directly tune condensate properties. Loss of RSKS-1 partially restores reproductive capacity, potentially by altering the extent or dynamics of GTBP-1 granule assembly and rebalancing stress-adaptive responses. These findings extend prior studies of mammalian S6K in oxidative stress responses and provide the first organismal evidence linking S6K activity to germline temperature resilience^42^.

In conclusion, these findings highlight phase-separated condensates as critical mediators of temperature-dependent reproductive fitness and establish GTBP-1 as key nodes linking environmental stress to germline continuity. Future studies should define the germline protein and RNA interactome of GTBP-1, and determine whether the internal partitioning of these interactors in germline stress granules represents a conserved organizational principle across species.

## Materials and methods

### Strains

The *C. elegans* strains used in this work are listed in **Table S1**. Animals were maintained on Nematode Growth Medium (NGM) plates seeded with *E. coli* OP50. All strains were grown at 20 °C unless otherwise specified.

### Construction of deletion mutants and transgenic strains

Deletion mutants and transgenic *C. elegans* strains were generated using the CRISPR/Cas9 system. For transgenic strains, coding sequences for GFP::3xFLAG or mCherry fused to a linker (GGAGGTGGAGGTGGAGCT) were inserted either upstream of the stop codon or downstream of the start codon of the target gene. For deletion mutants, sgRNAs were designed to target the coding sequence of the gene of interest. The sgRNA sequences used are listed in **Table S2**. Repair plasmids were constructed using the ClonExpress MultiS One Step Cloning Kit (C113-02, Vazyme), and plasmids for injection were prepared using the AxyPrep Plasmid Miniprep Kit (Axygen). The injection mixture contained 50 ng/μl Cas9 vector (pDD162), 50LJng/μl repair plasmid (for knock-in) or no repair template (for deletions), 5 ng/μl co-injection marker plasmid (pCFJ90), and 30 ng/μl each of 2–3 sgRNA plasmids, and was injected into young adult worms. Mutants and knock-in transgenic lines were confirmed by PCR amplification and DNA sequencing, and transgenic lines were further validated by fluorescence microscopy. Sequence of repair plasmids primers are listed in **Table S3**.

### Forward genetic screens

We used forward genetic screens to isolate *gtbp-1* suppressors. Basically, the *gtbp-1(ust600)* mutant animals (P0) were synchronized at the late L4 larval stage, collected in 4 mL M9 buffer, and incubated with 50 mM ethyl methanesulfonate (EMS) for 4 hours at room temperature with constant rotation. Animals were then washed with M9 three times and cultured under standard conditions. After 20 hours, adult animals were bleached. Eggs (F1) were distributed and raised on ∼100 9-cm NGM plates, resulting in a total of 30–50 plates for the F1 generation, each containing 50 to 100 eggs. These F1 animals were cultured until gravid, then subjected to bleaching to synchronize the F2 generation. The resulting F2 embryos were distributed onto new plates and incubated at 25 °C. Putative suppressor mutants were identified in the F3 generation based on their ability to produce viable and fertile progeny at the restrictive temperature of 25 °C, a phenotype not observed in the original *gtbp-1(ust600)* mutant. F3 animals were individually cloned, and their fertility was further confirmed in the F4 generation to ensure heritability of the suppressor phenotype. The causative mutation was identified through whole-genome sequencing and subsequently confirmed by the acquirement of multiple alleles.

### Brood size

L4 worms were placed individually onto fresh NGM plates. The numbers of progeny that reached the L2 or L3 stage were scored. A minimum of 15 animals per strain per condition were analyzed.

### Heat stress assays

For chronic mild heat stress: synchronized animals were continuously cultivated at the restrictive temperature of 25 °C from the L1 larval stage. Worms at the young adult stage were photographed for subsequent phenotypic analysis. For acute heat shock: synchronized L4 to young adult-stage worms were individually picked onto fresh NGM plates. The plates were sealed and subjected to a heat shock at 37 °C for 30 minutes. For recovery experiments, after heat shock plates were immediately returned to 20 °C for a specified period before phenotypic assessment or imaging.

### Microscopy and image analysis

For imaging of live adult animals, worms were immobilized in ddHLJO containing 0.25 M sodium azide and mounted on freshly prepared 2% agarose pads. For high-resolution imaging of embryos and germ cells within adult animals, worms were dissected in 2 µL of 0.4× M9 buffer containing 0.1 M sodium azide, and the released gonads or embryos were transferred onto freshly prepared 1.1% agarose pads. The Leica TNUNDER Imaging System was used, equipped with a K5 sCMOS microscope camera and an HC PL APO 100x/1.40-0.70 oil objective. Images were taken and deconvoluted using Leica Application Suite X software (version 3.7.4.23463). Identical microscope and camera settings were maintained for all comparative quantifications within a given experiment.

Stress granules were quantified within the meiotic region of the germline. ROIs of approximately 200 μm² were selected for analysis. For each condition, a minimum of 10 independent animals were analyzed, with 1–2 unit areas assessed per animal. The number of stress granules per unit area was manually counted and analyzed using ImageJ software.

The degree of colocalization between different fluorescently labelled proteins in germ cells was calculated using the Coloc2 plugin from ImageJ. All representative images are from at least three independent biological replicates, with a minimum of 10 animals or germlines analyzed per condition for quantification.

### RNAi screening

RNAi experiments were performed by feeding worms with *E. coli* HT115(DE3) expressing double-stranded RNA targeting specific genes. The RNAi bacterial clones were obtained from the Ahringer feeding library and confirmed by sanger sequencing. For analysis of stress granule dynamics, synchronized L1 larvae were raised on RNAi plates at 20 °C until the L4 or young adult stage, and then subjected to heat shock at 37 °C for 30 minutes. For brood size analysis, synchronized L1 larvae were raised on RNAi plates and maintained at the restrictive temperature of 25 °C throughout development until progeny counting. In all experiments, worms fed with the empty vector L4440 were used as the RNAi control. A list of the genes used in feeding RNAi experiments is provided in **Table S4**.

### Statistical analysis

Statistical analysis was performed using OriginPro 2017 (version 9.4.0). Details on the statistical test, the sample, and experiment number, as well as the meaning of error bars, are provided for each experiment in the corresponding figure legend, in the results and/or in the method details. Significance was defined as: *p<0.05; **p<0.01; ***p<0.001; ns, not significant, p>0.05.

## Acknowledgments

We are grateful to the members of the Guang laboratory for their comments. We are grateful to the International *C. elegans* Gene Knockout Consortium and the National Bioresource Project for providing the strains. Some strains were provided by the CGC, which is funded by the NIH Office of Research Infrastructure Programs (P40 OD010440).

## Author contributions

S.G., X.F., Z.Z., and D.C. conceptualized and designed the research; D.Z., Y.J., X.H., Y.K., J.C., W.W., D.X., and C.Z. performed the research; S.G. and D.Z. wrote the paper.

---

**Table S1. *C. elegans* strains used in this study.**

**Table S2. Sequences of the sgRNAs.**

**Table S3. Sequence of repair plasmids primers.**

**Table S4. List of genes used in the RNAi screening.**

